# Mapping heterogeneity drivers in human PSC-derived limbal stem cell differentiation through single-cell multi-modal analysis

**DOI:** 10.64898/2026.09.16.751974

**Authors:** J.A. Arts, D. Lima Cunha, S. Harjuntausta, M. Vattulainen, JGA. Smits, I. Bonthuis, D. Fragoso Bento, F. Bocci, H. Skottman, H. Zhou

**Affiliations:** Department of Molecular Developmental Biology, Radboud Institute for Molecular Life Sciences (RIMLS), Nijmegen, The Netherlands; Department of Human Genetics, Radboud University Medical Centre, Nijmegen, The Netherlands; Faculty of Medicine and Health Technology, Tampere University, Tampere, Finland; Norwegian Centre for Molecular Biosciences and Medicine (NCMBM), Nordic EMBL Partnership, University of Oslo, Oslo, Norway

**Author notes:** These authors contributed equally to the manuscript (J.A. Arts, D. Lima Cunha). These authors are co-corresponding authors (H. Skottman, H. Zhou).

## Abstract

Understanding the molecular underpinnings of stem cell differentiation is pivotal for generating appropriate cell types in cell-based therapies. Differentiation of human pluripotent stem cells (PSC) into corneal limbal stem cells offers a promising avenue to regenerate the corneal epithelium. However, current differentiation strategies remain inconsistent in efficiency and yield heterogeneous cell populations that incompletely recapitulate the regenerative properties of donor-derived limbal stem cells.

Here, we mapped the molecular landscape of cell states throughout the differentiation process. First, we construct PSC differentiation paths from single-cell RNA sequencing (scRNA-seq) data using the computational framework of optimal transport, identifying on- and off-track cell states toward the limbal stem cell state. Single cell Assay for Transposase-Accessible Chromatin sequencing (scATAC-seq) was performed to profile accessible genomic regions, and subsequently integrated with scRNA-seq data through gene regulatory network analysis to identify key drivers governing the diverse cell states. We showed that genomic enhancers play a major role in cell state determinations. Through this single-cell multi-modal approach, we identified potential transcription factors driving limbal epithelial lineage specification and off-track cell populations. Our findings provide a framework for rational optimization of PSC-derived limbal stem cell generation to advance the development of corneal cell therapies.

## Introduction

Understanding the precise control of cell fate decisions during stem cell differentiation is essential for generating appropriate cell types for cell-based therapies. In the core of cell fate control mechanisms are transcription factors (TF) and the chromatin environment, where TFs bind and enable contacts between gene promoters and genomic enhancers. These are key for proper gene expression and cell commitment [1, 2]. Thus, insights into the molecular mechanisms orchestrated by key TFs and the chromatin landscape during *in vitro* differentiation provide guidance to improve differentiation efficiency and consistency, as well as reducing unwanted cell populations in the cultures.

Human donor derived limbal stem cells (LSC) are currently used in cell-based corneal therapies, given their capabilities in homeostatic renewal to proliferate and repair the corneal epithelium [3]. However, this approach is not ideal for many reasons, e.g., unavailability of suitable donor tissue for autografting, especially in the case of bilateral limbal stem cell deficiency (LSCD), whereas allogeneic grafts often require long-term immunosuppression to avoid rejection. In this regard, differentiation of human pluripotent stem cells (hPSC) into corneal limbal stem cells, hereafter termed as induced LSC (iLSC), holds significant promise for cell-based corneal therapies, as evidenced by recently initiated clinical trials [4]. Human iLSC offer several major advantages in cell therapy such as easy accessibility, abundance and the possibility of personalized autologous treatments, the latter bypassing the need for immune suppression. More recently, “off-the-shelf” allogeneic hPSC-derived cells also became an attractive option, allowing for faster, more cost-effective, and scalable treatment options [5].

Up to date, multiple iLSC differentiation methods have been developed (reviewed in [6]). Despite significant progress in developing differentiation protocols, achieving robust and reproducible iLSC population still requires substantial improvement [7]. Previously, our single-cell RNA sequencing (scRNA-seq) analysis on differentiating iLSC derived from several hPSC lines showed multiple cell states in cultures at the end of differentiation, not only intended iLSC, but also mesenchymal and pluripotent-like cells, both of which were detected transiently and persistently across the differentiation process [8]. Importantly, this heterogeneity seemed to be PSC line-dependent, which is a common problem in PSC differentiation[7]. Understanding the relevance of the diverse cell states and mechanisms behind line-dependent variability in the differentiation process is essential for further optimization of the method and advance future applications of PSC-derived cell therapy technology.

In the previous study, pseudotime analysis was performed to provide initial insights into transcriptional changes among these cell states along differentiation [8]. However, pseudotime-based methods do not take advantage of the temporal information in our time course differentiation data, and more importantly, do not explicitly connect progenitor states to their descendants [8]. As a result, it remains unclear how cell states at early time points give rise to the later ones, and accordingly, it is yet to be determined whether a certain cell state is on-track or off-track along the desired iLSC trajectory. To address these questions, optimal transport (OT)-based computational frameworks represent cell state transition processes as probabilistic flows from early to late time points [9, 10]. Specifically, Waddington OT (WOT) [11] applies OT modeling on time-series single-cell data to reconstruct developmental trajectories, thereby extending Waddington’s classical “epigenetic landscape” concept [12]. In essence, WOT leverages cell-cell similarity, temporal information and estimated rates for cell growth and apoptosis to identify ancestor and descendant relations between cells along a time course. By establishing quantitative links between ancestor and descendant cell states, WOT analysis may reveal previously unresolved cell state trajectories, and identify on-track or off-track cell states towards iLSC.

Importantly, the control mechanisms of on-track or off-track cell states, including key TFs and the chromatin environments where these TF function, is completely unexplored in iLSC differentiation. To this end, single-cell Assay for Transposase-Accessible Chromatin sequencing (scATAC-seq) is informative and complementary to scRNA-seq data, as scATAC-seq profiles accessible chromatin regions, including promoters and genomic enhancers where TFs bind and regulate downstream gene expression [13, 14]. Integrating scRNA-seq and scATAC-seq data by constructing gene regulatory networks (GRN), captures the potential causal and combinatorial TF and target gene relationships underlying transcriptional regulation, predicting which key TFs drive specific cell states. GRN-based methods can more reliably predict TFs than using gene expression data or chromatin accessibility data with TF motif detection alone. scANANSE, a multi-modal based tool developed by our group [15], utilizes integrated GRN to calculate TF influence scores, enabling systematic ranking of key TF between cell states.

In this study, we applied WOT to previously generated scRNA-seq data on iLSC differentiation (Figure 1A) to determine the differentiation trajectory and identify which cell states contribute, or not, towards the iLSC state in individual PSC lines [8]. Furthermore, we performed scATAC-seq (Figure 1A) to profile accessible regions in the diverse cell states detected across the differentiation process. By integrating scRNA-seq and scATAC-seq datasets, we constructed GRNs using scANANSE and identified TFs key to cell specification in the diverse on-track and off-track cell states (Figure 1B). Finally, we compared molecular signatures between specific cell states to dissect the mechanisms governing each state in a PSC line-dependent manner. This study provides insights into the regulatory landscape governing iLSC differentiation, suggesting potential strategies to enhance differentiation efficiency.

**Figure 1:**
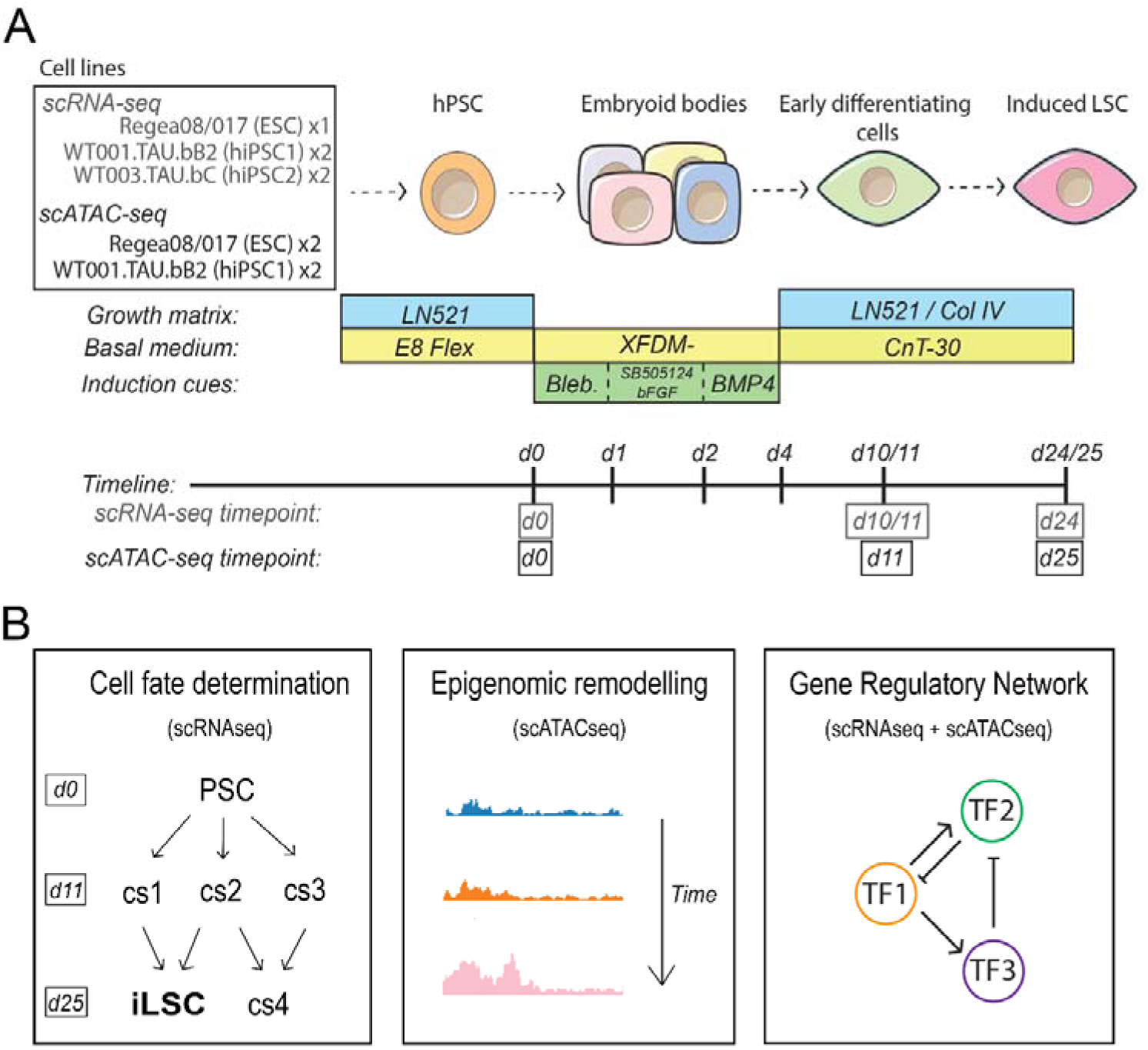
iLSC differentiation protocol and schematic overview of the study. A) Differentiation protocol and cell line composition for scRNA-seq [8] and scATAC-seq (performed in this study) (figure modified from Vattulainen et al., (2024) [8]. B) Schematic overview of the study, depicting the key analyses performed with scRNA-seq and scATAC-seq datasets. Abbreviations: Bleb, blebbistatin; LN521, laminin521; PSC, pluripotent stem cells; cs, cell state; iLSC, induced limbal stem cell; TF, transcription factor.

## Results

### Cell state trajectories in differentiation of PSCs to iLSCs

We previously performed a scRNA-seq time course study to characterize differentiation of human PSCs to iLSC using an ESC line (hESC) and two iPSC (hiPSC1 and hiPSC2) lines, showing cellular heterogeneity and variable iLSC differentiation efficiency [8]. In that study, the main observed cell states included pluripotent stem cells (PSC) at the start of differentiation (day 0), a predominant epithelial cell cluster (early epithelial, EE) and three smaller clusters with mesodermal signatures (mesoderm-like (ML) 1, 2 and 3) that emerged during differentiation (day 10/11), and two epithelial cell states appearing at the end of differentiation (day 24): the intended iLSC and an extra state annotated as late epithelial cells (late-epi). Additionally, a population of cells with PSC-like characteristics were detected on day 10/11 and more prominently on day 24 (Figure 2A). The cellular heterogeneity was apparently different between PSC lines, with highest percentage of EE and iLSC derived from the hESC line, the lowest from hiPSC2 and intermediate from hiPSC1 differentiations. Importantly, PSC-like cells were predominantly observed in differentiation from the hiPSC1 and hiPSC2 lines, and only rarely from the hESC cell line, suggesting line-dependent differences linked to differentiation efficiencies [8].

**Figure 2:**
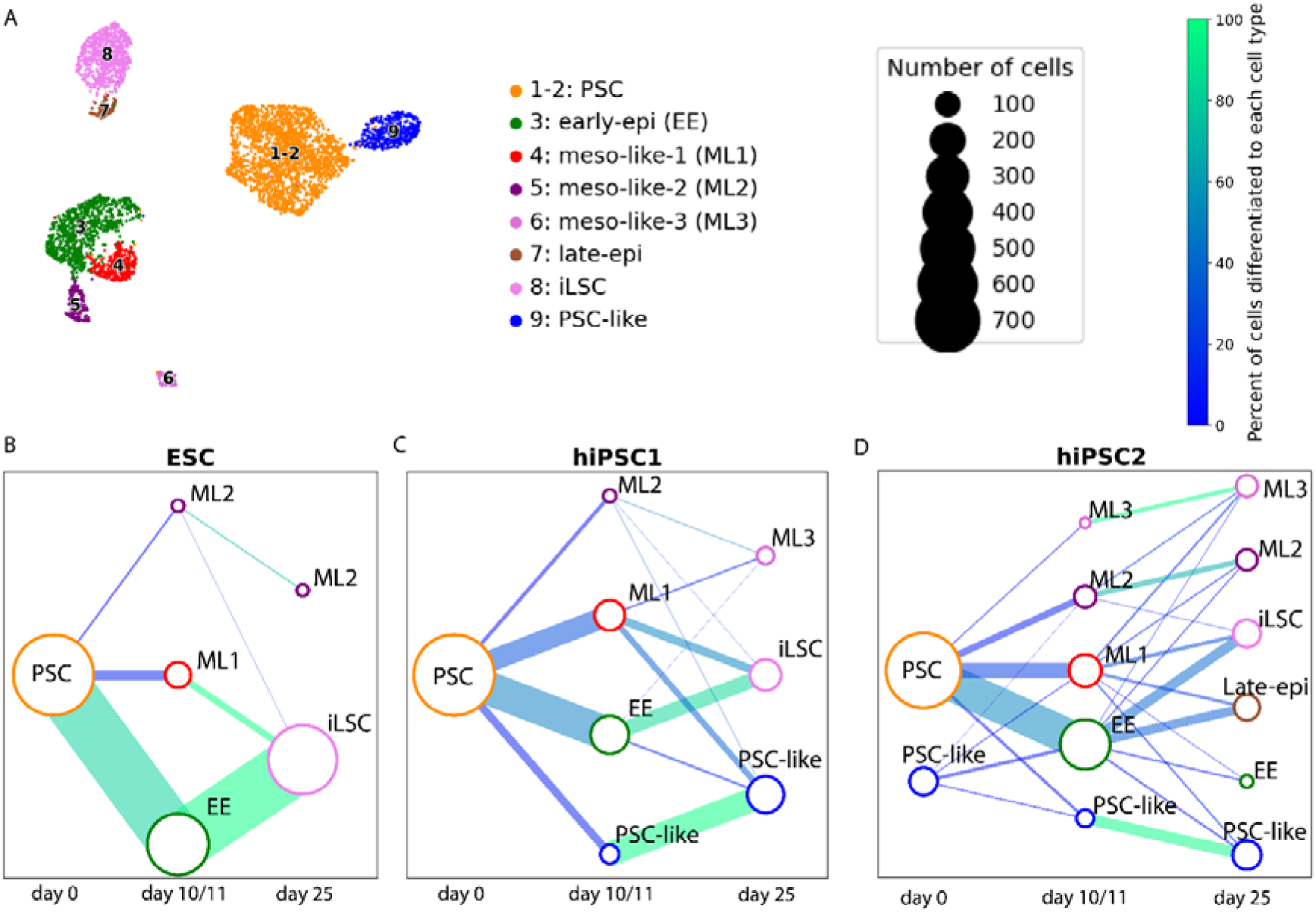
Predicting ancestors and descendant scRNA-seq cell states through WOT. A) Adapted scRNA-seq UMAP from [8]. B-D) Networks depicting ancestors and descendant cell states (>10 cells) comparing day 0 against day10/11 and day10/11 against day 24 across cell lines hESC (B), hiPSC1 (C), hiPSC2 (D). Node (dot) sizes indicate the absolute number of cells for each scRNA-seq cell state (Supplementary Table 1) at a specific timepoint. The color of the edges between nodes indicate percentages of cells from a single cell state at a previous timepoint transitioning into cell state states at the subsequent timepoint. Edge (line) width corresponds to the total number of transitioning cells from all cell states at one timepoint into the subsequent timepoint.

To examine the mechanism behind PSC line dependent differentiation heterogeneity, scRNA-seq data was reanalyzed using WOT to track ancestor-descendant cell trajectories during iLSC differentiation (Figure 2A, Supplementary Table 1) [8]. Transition probabilities between cell states of the adjacent time points were calculated (Supplementary Figure 1) and visualized as networks between these time points (Figure 2B-D). To gain more insight into cell fate commitment at the level of individual cells, we further computed cell fate scores (0–1) that indicate the probability of individual cells to progress into the adjacent later cell states (Supplementary Figure 2; Supplementary Table 2). Analyses compared day 0 vs 10/11 and day 10/11 vs 24, and were performed separately for each PSC line to detect line-dependent differences (Figure 2B-D, Supplementary Figure 1; Supplementary Table 2).

Confirming our previous findings, the hESC line was the most efficient in generating iLSC, with >95% cells consisting of iLSC on day 24 (Figure 2B; Supplementary Table 1). Among PSC transitioning from day 0 to day 10/11, WOT predicted 82% and 14% of PSC became EE and ML1, respectively, and a high percentage of WOT cell fate scores above 0.5 (99%) for these transitioning cells, indicating this trajectory has a high probability (Supplementary Table 2). In contrast, only 3% of cells were transitioning to ML2, with a very low cell fate score of 0.001, indicating that hESC is not likely differentiated to ML2. Among cells transitioning from day 10/11 to day 24, iLSC were predicted to derive mostly from EE and ML1 (84% and 10% of all day 10/11 cells, respectively). This trajectory has a high percentage of cells with WOT cell fate scores above 0.5 (98%), indicating that EE and ML1 are likely the intermediate cell states on-track towards the iLSC state. In contrast, <1% of the cells was transitioning from ML2 to iLSC, and 2/3 of ML2 continued as ML2 onto day 24 (Figure 2B, Supplementary Figure 1). This suggests that ML2 may be a small off-track population, with no relevant contribution to the iLSC state.

In hiPSC1, WOT predicted more complex trajectories, consistent with the higher cellular heterogeneity and lower iLSC yield (35% of total cells on day 24) observed previously [8] (Supplementary Table 1). On day 0, 99.7% of cells were annotated as PSC and 0.3% as PSC-like (Supplementary Table 1). Similarly to the hESC line, PSC predominantly gave rise to EE and ML1 states by day 10/11 (48% and 31% of all transitioning cells, respectively), with a high percentage of cells with cell fate scores above 0.5 (85%) (Figure 2C, Supplementary Figure 1, Supplementary Table 2). Transitioning cells between day 0 to day 10/11 showed higher percentage to ML2/ML3 and PSC-like, as compared to hESC. Among these, 8% became ML2/ML3 of day 10/11 cells, whereas 12% became PSC-like cells by day 10/11. Among day 24 cells, iLSC were shown to mainly derive from EE and ML1 of day 10/11 cells (22% and 12%, respectively), with a high percentage of cells with cell fate scores above 0.5 (72%) (Figure 2C, Supplementary Figure 2, Supplementary Table 2), although these transitioning cells have lower percentage and lower cell fate scores as compare to those of hESC. Of note, a high percentage of PSC-like (34%) stayed as PSC-like cells during the differentiation from day 10/11 to day 24 (Figure 2C, Supplementary Figure 1).

In hiPSC2 differentiation, WOT prediction showed the most complex trajectory patterns, corroborating with the highest cell state heterogeneity and, consequently, the lowest iLSC yield (22% of all day 24 cells) among the three PSC lines reported previously (Figure 2D, Supplementary Table 1). On day 0, 88% of cells were annotated as PSC, and already 12% as PSC-like (Supplementary Table 1). Similar to both hESC and hiPSC1, day 0 PSC predominantly gave rise to day 10/11 EE and ML1 states (Figure 2B–D) (51% and 21% of all transitioning cells, respectively), with a high percentage of cells with cell fate scores above 0.5 (93%) (Supplementary Figure 1, Supplementary Table 2). A small proportion of day 0 cells also transitioned into ML2 (10%), ML3 (2%), or PSC-like (4%) states on day 10/11 (Figure 2D). Among all cells transitioning from day 10/11 to day 24, 15% were from EE to iLSC, lowest among all three lines, whereas 13% were from EE to late-epi, highest among the three lines. PSC-like, ML2, and ML3 states largely originated from their corresponding cell states from day 10/11 (Figure 2D, Supplementary Figure 1, Supplementary Table 2).

Overall, while confirming previously reported cell line-dependent heterogeneity during iLSC differentiation, our WOT trajectory inference analysis predicted EE and ML1 as cell states being on-track to iLSC. In contrast, ML2 and ML3 seemed to be off-track cell states in the differentiation process. Likewise, PSC-like cells were determined as persistent and off-track cells, mainly present in differentiation of the hiPSC lines but not the hESC line.

### Chromatin accessibility-based cell state characterization

To further dissect the molecular signatures and the mechanisms driving diverse cell states throughout iLSC differentiation, we performed scATAC-seq to capture chromatin accessibility profiles to complement the transcriptomic data. Here we focused on hESC and hiPSC1 that differentiated to iLSC more robustly, as compared to hiPSC2. We differentiated these lines toward iLSC, following the same protocol as used for scRNA-seq (Figure 1A) [8], and collected samples from two independent experimental replicates at corresponding time points (days 0, 11, and 25).

From the two scATAC-seq replicas a total of 7452 cells passed quality control using snapATAC2 [16]. We then applied the batch-correction method Harmony [17] to integrate the datasets for downstream cell state annotation. Finally, we performed Leiden clustering[18] on open chromatin profiles from scATAC-seq, identifying seven clusters [19] (Figure 3A & Supplementary Table 3).

**Figure 3:**
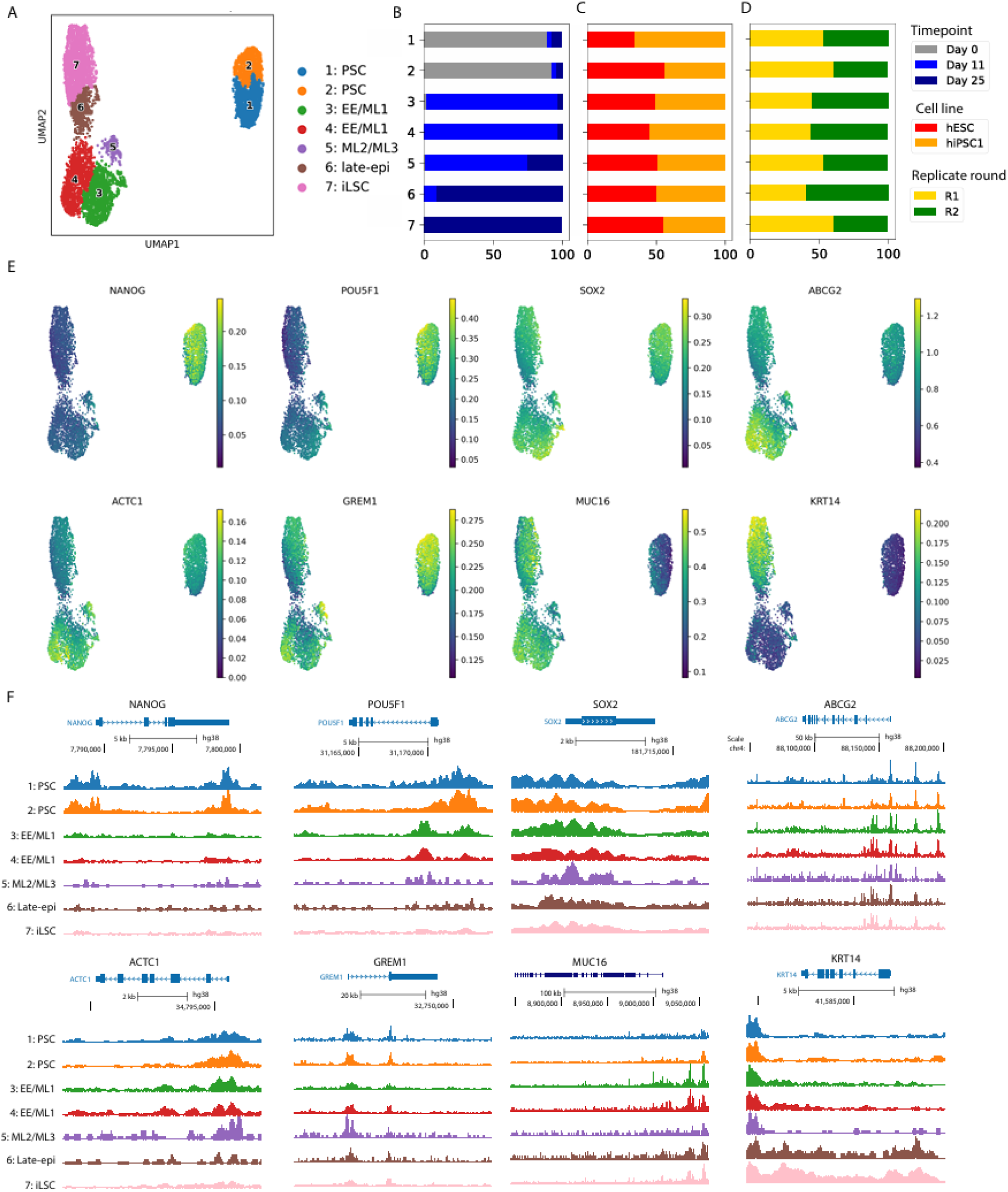
Characterization of cell states in scATAC-seq. A) Clusters and cell states identified in scATAC-seq. B-D) Proportions of time points (B), cell lines (C) and replicas (D) per cluster. E-F) Pseudobulk expression of genomic accessibility (E) and UCSC genome browser tracks (F) of NANOG, POU5F1, SOX2, ABCG2, ACTC1, GREM1, MUC16 and KRT14. *PSC = pluripotent stem cell, EE = early-epi, ML = meso-like, iLSC = induced limbal stem cell.*

Generally, scATAC-seq clusters showed minor variations (percentages ranging mostly between 40 and 60%) between the two replicas and the two PSC lines (Figure 3B-D, Supplementary Table 3). Although both PSC lines contributed to all clusters, 2x2 Fisher-style permutation test showed significant differences in the cell proportions of three clusters (1, 2 and 7) between hESC and hiPSC1 lines (Supplementary Table 4). To assign cell state identities to the scATAC-seq clusters, we applied label transfer with scANVI [20] (Supplementary Table 5), integrating chromatin accessibility with gene expression profiles by matching accessible genomic regions detected in scATAC-seq to nearby genes detected in scRNA-seq. This enabled the annotation of scATAC-seq clusters to the corresponding cell states defined in the scRNA-seq dataset.

Clusters 1 (1,374 cells) and 2 (828 cells) containing mostly day 0 cells (Figure 3B) were predicted to associated mainly with PSC (Supplementary Table 5). Both clusters displayed high chromatin accessibility around canonical pluripotency genes, including *SOX2*, *NANOG*, *POU5F1*, and *MT1G* (Figure 3E and F, Supplementary Figure 3), consistent with PSC identity. Given their similar accessibility profiles and enrichment for PSC-associated annotations, clusters 1 and 2 were merged and annotated as PSC. However, approx. 10% and 7% of cells in clusters 1 and 2, respectively, came from day 11 and day 24 samples, which we suspect may be PSC-like cells (Figure 3B and Supplementary Table 3).

Clusters 3 (1,252 cells) and 4 (1,223 cells) comprised mostly day 10/11 cells (Figure 3B) and were predicted to correspond to EE and ML1 states (Supplementary Table 5). Both clusters shared open chromatin accessibility at loci of many EE and ML1 genes validated in our previous study [8], e.g., *ABCG2* and *ACTC1* (Figure 3E and F), associated with LSC and mesodermal differentiation, respectively [21, 22]. Unlike the transcriptomics data, clusters 3 and 4 lacked distinct separation in chromatin accessibility features, and therefore were merged and annotated as EE/ML1.

Cluster 5 (221 cells) contained a mix of day 10/11 (75%) and day 25 cells (25%; Figure 3B). This cluster had the highest proportion of predicted ML2 and ML3 states (Supplementary Table 5), exhibiting high chromatin accessibility at *ZEB2* and *COL1A1* loci (Supplementary Figure 3B) as well as for GREM1 (Figure 3E and F), a gene known to induce mesenchymal gene expression [23]. These three genes were previously described as markers of ML2 and/or ML3 states [8] and therefore annotated as ML2/ML3.

Clusters 6 (709 cells) and 7 (1,845 cells) were composed primarily of day 25 cells (Figure 3B). Both clusters showed high proportions of predicted late-epi and iLSC states, with cluster 6 enriched for late-epi and cluster 7 for iLSC states (Supplementary Table 5). Although minor, chromatin accessibility differences between these two clusters were noticeable. Cluster 6 retained higher accessibility signals at *MUC16* and had lower at LSC-associated gene *KRT14* than cluster 7, and vice versa (Figure 3F, Supplementary Figure 3D). Thus, based on the correlations with gene expression pattern previously observed in scRNA-seq cell states [8], cluster 6 was annotated as late-epi and cluster 7 as iLSC.

Together, these findings demonstrate that chromatin accessibility profiles recapitulate the major cell states identified by scRNA-seq and show that progressive epigenomic remodelling occurs during iLSC differentiation.

### Unique accessible chromatin regions in genomic enhancers control cell state specific genes during iLSC differentiation

Previous studies have shown that key accessible regions defining cell state identity are often found in enhancer regions rather than in promoters [24–26]. Therefore, we determined the genomic locations of accessible regions identified in different stages during iLSC differentiation. We first associated identified peaks within a ±2 kb window of a gene body defined as promoter regions [27]. Subsequently, we linked the remaining peaks to genes within a ±50 kb window defined as putative enhancer regions, as previously described [1]. We identified a total of 440,306 peaks across all cell states, including PSC (282,288), EE/ML1 (260,838), ML2/ML3 (73,044), late-epi (76,985), and iLSC (228,126) (Supplementary Table 6). Approximately 70-86% identified peaks associated with different cell states were located in enhancer regions (Supplementary Table 6).

Next, we questioned whether unique enhancers to a given cell state would play a more determinant role in defining that corresponding cell state than promoters. To address this, we defined unique peaks as those present in only one cell state but absent in all others. The PSC state showed the highest number of unique peaks and the highest ratio of unique peaks to all peaks annotated to PSC, with 44% (124,080) of all peaks mapped to PSC. EE/ML1 and iLSC states had intermediate number and ratio, 45,088 (17% of those mapped to EE/ML1) and 42,670 (19% of those mapped to iLSC), respectively. ML2/ML3 and late-epi had the lowest number and lowest ratio, 5,000 (0.07% of those mapped to ML2/3) and 1,483 (0,02% of those mapped to late-epi), respectively (Supplementary Table 7). Similar to all identified peaks, we examined whether these unique peaks are located in enhancer or promoter regions. As expected, for all cell states, unique peaks showed a large majority located in enhancer regions (95-97%), and were significantly enriched for enhancer/promoter ratio, as compared to all peaks (Figure 4A; Fisher’s exact test: OR = 3.592, p < 1 * 10^-6^).

**Figure 4:**
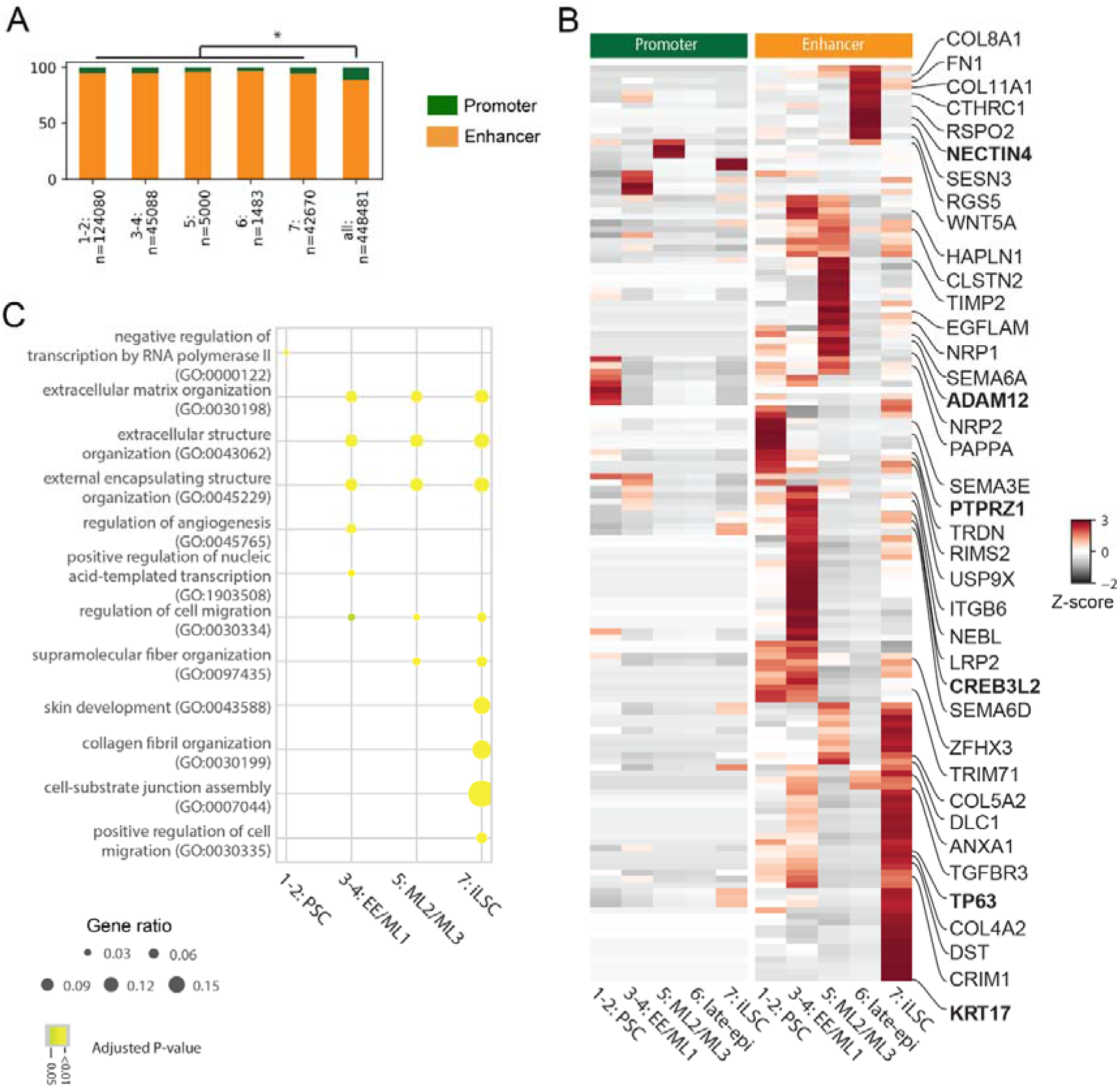
Biological processes identified for unique peaks linked to highly variable genes. A) Enhancer and promoter distribution for unique peaks for identified cell states merged clusters. * indicates a p-value <1e-6 (Fisher’s exact test). B) Heatmaps of scATAC-seq depicting Z-scores (enhancer-to-gene signal) from peaks linked to previously defined highly variable genes in scRNA-seq (log2FC>1.5 and p-adjusted value <0.05). The top 5 HVGs with highest peak signal at enhancers are located on the right of the plot. C) Top 10 merged GO-terms on enhancer-to-genes signal of previously identified HVGs (log2fc>1.5, *padj < 0.05) for unique peaks. Dot size indicates the ratio of genes found for a specific GO-term. The color of the dot indicates the adjusted p-value. PSC = pluripotent stem cells, EE/ML1 = early-epi/meso-like-1, ML2/ML3 = meso-like-2/meso-like-3, iLSC = induced limbal stem cells.*

As highly variable genes (HVGs) in scRNA-seq data typically represent the main differences between cell states, we examined whether unique peaks linked to HVGs are most variable between cell states and could drive the diverse cell state identify. In both promoters and enhancers, unique peak signals near HVGs were mostly distinct for each cell state, such as PTPRZ1 (PSC), CREB3L2 (EE and iLSC), ADAM12 (ML2/ML3), NECTIN4 (late-epi), TP63 (iLSC) and KRT17 (iLSC) (Figure 4B). In line with this, biological processes of HVGs linked to unique peaks in enhancers revealed more distinct patterns in cell states, shown by Gene Ontology (GO)-term enrichment analysis (Figure 4C). In contrast, much less distinct accessibility signals between cell states were detected in all peaks when they were mapped to HVGs (Supplementary Figure 4A). Consistent with this, GO annotation showed broad biological process terms from HVGs linked to all identified peaks (adjusted p-values < 0.05) but not specific for individual cell states (Supplementary Figure 4B). These observations demonstrate that unique enhancers play a major role in determining different cell states. Interestingly, there was little overlap between promoter- and enhancer-associated peaks linked to HVGs associated to cell states (Figure 4B), suggesting distinct roles of promoters and enhancers in driving expression of different genes.

### Identification of TFs driving cell states through gene regulatory network analysis

As TFs are key regulators by binding to promoters and enhancers to regulate target gene expression, thereby driving cell state determination, we sought to identify TFs governing cell state heterogeneity that arose during iLSC differentiation. We applied gene regulatory network (GRN)-based analysis that integrates activities of promoters, genomics enhancers and expression of their corresponding target genes. scANANSE [15] is such a method for single-cell datasets and ranks TFs in order of importance for different cell states. For each cell state, we compared its GRN against a reference network averaged from all cell states, and determined TF influence scores (scale 0–1) that represent the importance of TF for defining the cell states. As our focus here is on cell state-specific differences, for the robustness, we used combined scATAC-seq data of hESC and hiPSC1.

We generated pseudobulk from both modalities for scANANSE analysis and attempted to link them in a cell state specific manner. For late-epi and iLSC cell states, pseudobulk of both modalities were defined, and they could be directly coupled. However, for other cell states, the global scATAC-seq profiles were not distinguishable between similar cell states, e.g., those between PSC and PSC-like, or between EE and ML1, or between ML2/ML3. In these case, the GRNs were generated from their cell state-specific scRNA-seq pseudobulk and the combined scATAC-seq pseudobulk. For instance, the combined scATAC-seq pseudobulk of PSC (cluster 1 and 2) together with specific PSC and PSC-like scRNA-seq pseudobulk that distinguished well between these cell states was used to generate GRNs for PSC and PSC-like. Similarly, scATAC-seq pseudobulk of EE/ML1 was used for GRNs of EE and ML1, and that of ML2/ML3 was used for GRNs of ML2 and ML3.

Many TFs detected in GRNs with high influence scores in PSC and PSC-like were shared, such as SOX2, NANOG and POU5F1, ZIC2, ZIC3 and ZIC5 (Figure 5A, Supplementary Figure 5). Nevertheless, several TFs were detected to be PSC-like specific, including SP8, PAX5, NKX1-2 and PRDM14 (Figure 5A,B). In ML2 and ML3, TFs with high influence scores such as FOXC2, NR2F1, and ZEB1 were shared between these two states (Figure 5A,B). Although the chromatin accessibilities are largely similar between ML2 and ML3, some TFs showed influence score specificity for either, e.g, FOXC1 and MYBL1 for ML2 and HOXA13, HOXA10, GATA4 for ML3, highlighting distinctions between these cell states (Figure 5A).

**Figure 5:**
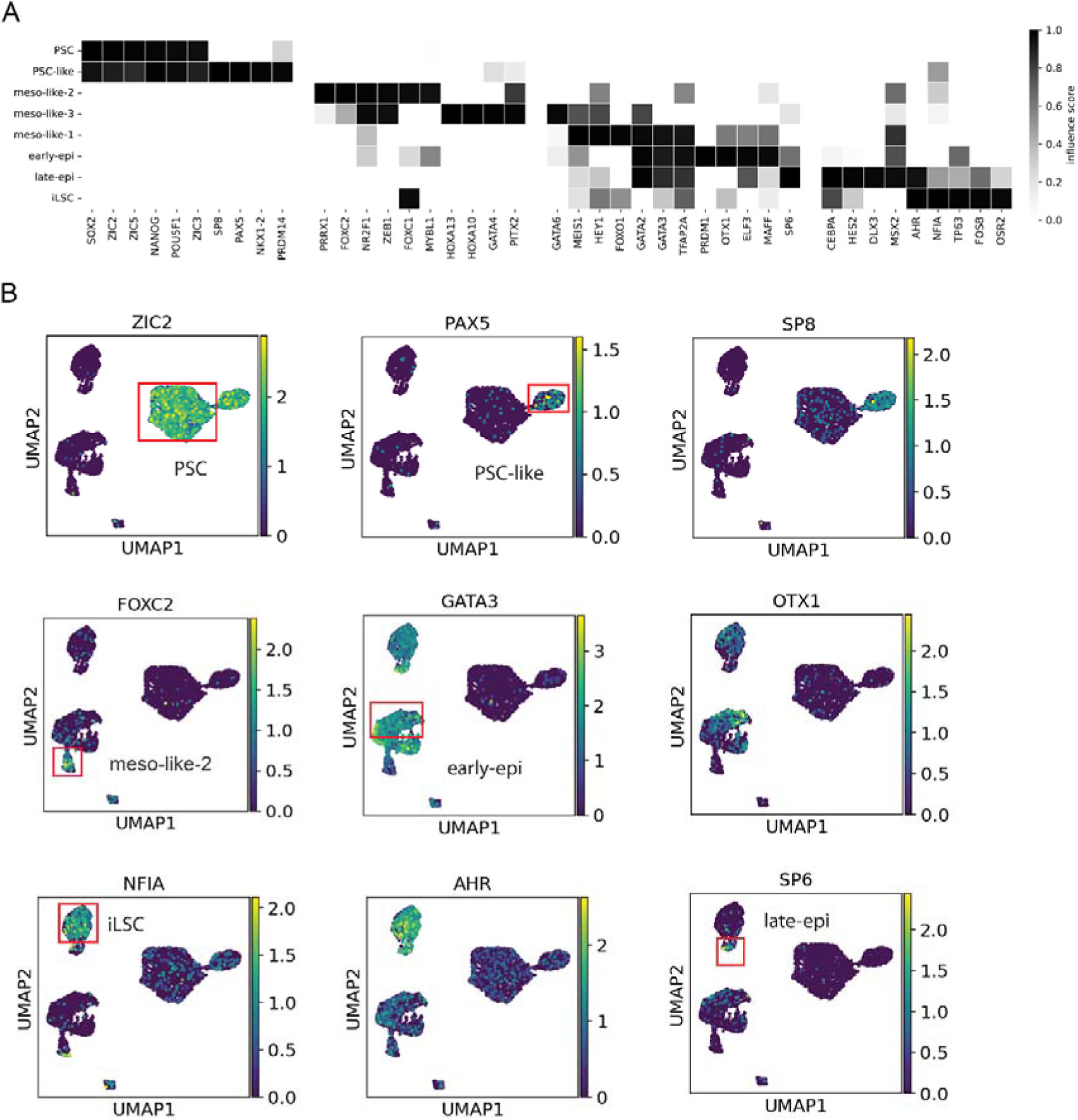
Identification of key TFs controlling cell states through scANANSE. A) Top 6 influential TFs per cell state identified through comparison of its cell state GRN to an average GRN constructed from all peaks. B) Normalized gene expression from scRNAseq dataset of distinct influential TFs: ZIC2 for PSC, PAX5 and SP8 for PSC-like, FOXC2 for ML2, GATA3 and OTX1 for EE/ML1, NFIA and AHR for iLSC, and lastly, AHR and SP6 for late-epi.

TFs with high influence scores in EE and ML1 were hardly specific for these two cell states. For example, MEIS2, HEY1 and GATA2 showed high influence scores not only in ML1 and EE, but also in ML2, ML3 and late-epi. GATA3, TFAP2A, OTX1, EFL3, MAFF had high scores in ML1, EE, late-epi and even medium scores in iLSC (Figure 5A, B). These observations suggest that ML1 and EE are intermediate states on-track towards iLSCs, consistent with the findings from WOT analysis.

For late-epi and iLSC states, TFs with high influence scores were mainly shared, e.g., CEBPA, HES2, AHR, reflecting the close relationship between these two states (Figure 5A,B). NFIA, TP63, FOSB showed relatively high scores in iLSCs. Interestingly, in addition to the high score in late-epi, MSX2 had also relatively high score in ML1, 2, 3 and EE (Figure 5A), indicating that the late-epi may either be a delayed on-track state towards iLSCs or a slightly different epithelial state closely related to iLSC. Finally, it is important to note that our GRN analysis identified a broad set of LSC-related TFs with high influence scores in the iLSC state, including TP63 [28], CEBPD, EHF[29], KLF3, KLF4, KLF6, EGR1, HES1 [30], FOSL2 [1], ELF1, RUNX1 and SMAD3 [31] (Supplementary Table 8), highlighting the LSC character of these cells.

### Enhanced expression and chromatin accessibility of TFs in persistent PSC-like cells

Our WOT analysis indicated that iLSC were mainly derived from on-track EE and ML1 cells (Figure 2), whereas PSC-like cells were cells that were persistently present at day 10/11 and day 24 of the differentiation process and not contributing towards the iLSC fate. As such, we speculated that differences between PSC and PSC-like are key to high iLSC differentiation efficiency. Therefore, we directly compared GRNs that contained integrated information of both gene expression and chromatin accessibilities between PSC and PSC-like cells to identify the differences between these cell states and the TFs driving the PSC-like identity.

A number of TFs showed high influence scores as well as increased gene expression in either cell states (Figure 6A). Among these TFs, OSR2, EBF1, HMX2, and POU3F2 showed both higher expression and influence scores, forming an inter-connected GRN in PSC cells (Figure 6A, Supplementary Figure 5A). In contrast, SP8, PAX5 and HOXB2 displayed higher expression and influence scores in PSC-like cells, and regulated each other within an inter-connected network (Figure 6A, Supplementary Figure 5B).

**Figure 6:**
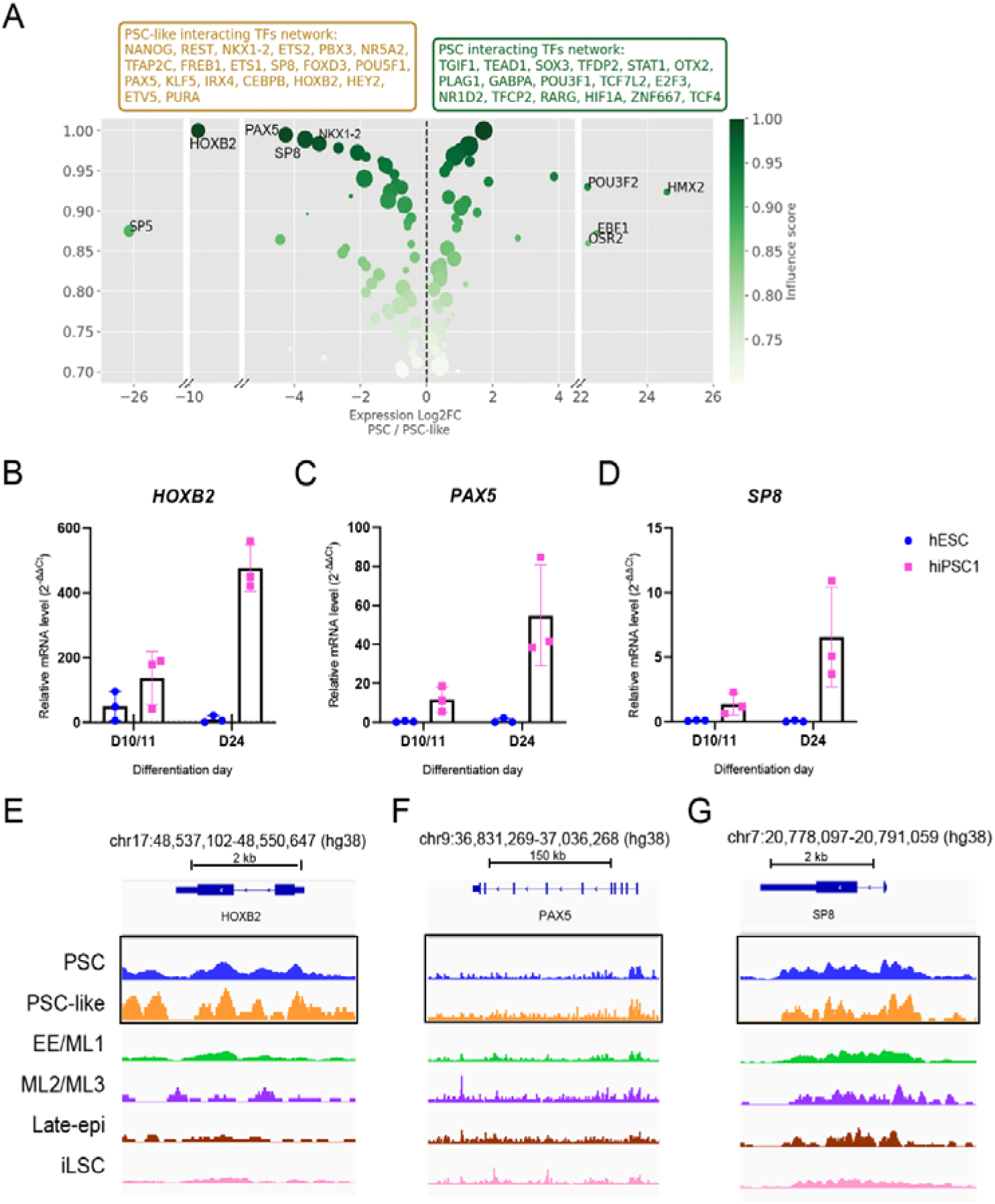
Top influential and differentially expressed factors in direct network comparisons of PSC and PSC-like. A) TFs identified through direct comparisons of GRNs constructed from unique peaks between PSC against PSC-like. The x-axis depicts the log2 foldchange between PSC over PSC-like (positive values, high in PSC; negative values, high in PSC-like). Negative values indicate an elevated log2 foldchange in PSC-like over PSC. The y-axis depicts scANANSE influence score of factors >0.7. B-D) Relative mRNA levels (qPCR) of HOXB2 (B), PAX5 (C) and SP8 (D) in ESC and hiPSC1. E-G) UCSC genome browser tracks for HOXB2 (E), PAX5 (F) and SP8 (G) loci. Tracks from PSC (blue) and PSC-like (orange) states were boxed for easier comparison.

Given that hiPSC1 showed a higher tendency than ESCs to develop PSC-like cells during differentiation, we further examined with RT-qPCR whether the three candidate TFs PAX5, SP8, and HOXB2 exhibited distinct expression patterns at the mRNA level between the two cell lines over time. Interestingly, although mRNA expression of these three genes were not measurable at the start of the differentiation, all three genes exhibited higher mRNA levels in hiPSC1 than in ESCs, and the disparities increased over time. At day 10/11 *PAX5, SP8*, and *HOXB2* expression was elevated in the hiPSC1 cell line relative to the ESC cell line (Figure 6B-D). At differentiation day 24, the differences at mRNA levels increased further in hiPSC1 compared to ESC (Figure 6B-D), suggesting that these TFs may contribute to the enhanced persistence of PSC-like states in hiPSC1. Corroborating with gene expression, chromatin accessibility of all 3 genes was markedly higher in PSC-like cells (cells in scATACseq PSC cluster that derived from day 11 and day 24 samples), compared to PSCs (day 0 cells in scATAC PSC cluster) (Figure 6E-G).

## Discussion

In this study, we performed integrative scRNA-seq and scATAC-seq analyses to investigate the molecular dynamics underlying iLSC differentiation. Through the WOT modeling approach, we characterized heterogeneous cell states during differentiation, with EE and ML1 on-track to iLSC, and PSC-like, ML2 and ML3 being likely off-track cell states. Chromatin accessibility scATAC-seq analysis appeared to be less distinct to detect cell states than scRNA-seq, but demonstrated that cell state determining regulatory regions are primarily located in enhancers rather than promoters. Gene regulatory network analysis integrating both gene expression and chromatin accessibility confirmed well-known LSC-associated TFs as key drivers of iLSC identity, including TP63, CEBPD, ATF3, FOSL2, RUNX1, and SMAD3. Importantly, we also identified PAX5, HOXB2 and SP8 as TFs governing PSC-like cells.

Understanding cell differentiation trajectories and recognizing on-track and off-track cell populations are important for optimization of differentiation cultures and future cell therapy applications. For example, the on-track cell states can be stimulated and off-track populations can be suppressed or even eliminated via modifying culture conditions. In our previous study, we applied pseudotime analysis to delineate the diverse cell populations arisen during differentiation [8]. However, pseudotime methods are based on clustering similar patterns in high-dimensional space to predict the order of appearing cell states, but cannot reliably predict the origin and differentiation destiny of any given cell state. Therefore, in this study, we applied WOT analysis that integrates cell-cell similarity with temporal information. WOT predicted that, among cell states appearing at the intermediate stage (day10/11), EE and ML1 were on-track states towards iLSCs, whereas PSC-like, ML2, 3 were off-track states further persisting at later time points. Additionally, the previous pseudotime analysis predicted trajectories such as EE differentiating into ML2 and subsequently into ML3, whereas WOT showed that ML2 and ML3 were unlikely originated from EE, but were cell populations emerging at the intermediate and persisting till the final stage. WOT analysis directly linked EE and ML1 to the iLSC state, providing new insights into the lineage commitment during differentiation [8]. It is important to note that the off-track cell states were mainly observed in hiPSC1 and hiPSC2 that were overall less efficient in iLSC generation and predicted by WOT to have more complex differentiation trajectories than hESC. Our analysis using WOT on each individual line illustrated a clear ancestor and descendant relationship of the complex cell states in these lines.

To further dissect the control mechanisms of diverse cell populations, we performed scATAC-seq for chromatin accessibility, to corroborate scRNA-seq gene expression data. Apparently, chromatin accessibility between the diverse cell states were much less distinct than gene expression between cell states, an observation that has been reported previously [32]. As expected, the PSC state was the most distinct from the rest, with the others relatively more similar between each other. Furthermore, in the analysis of unique peaks, we also observed a large difference between the ratio of unique peaks to all peaks annotated to a specific cell states. In PSC, 44% of peaks were unique to this state, highest among all cell states, whereas in late-epi, the least unique cell state, only 0.02% peaks were unique. Importantly, through unique peak analysis, we showed that peaks located in enhancer regions were more meaningful in defining cell states, than those in promoter regions, shown by their nearby HVGs and the gene functions specific to the cell states. This is in agreement with previous reports, showing that enhancers play a major role in defining cell type–specific gene expression [33, 34].

By integrating single-cell gene expression and chromatin accessibility, GRN analysis with scANANSE enabled us to identify TFs not only based on their expression levels, their binding to enhancers and promoters, but also their regulatory influence on downstream target genes. Multiple canonical TFs such as CREPA, TP63 and FOSB associated with LSC maintenance were identified in iLSC with high influence scores. Many of these were shared between iLSC and late-epi states, among which TP63, which had slightly higher influence score in the iLSC state, consistent with its role in corneal/limbal stem cells[28]. Late-epi shared high similarity with iLSC, but also shared multiple influential TF with iLSC precursor state EE/ML1. This is consistent with pseudotime analysis of scRNAseq data in our previous study, suggesting this cell state being either delayed iLSC or stuck in differentiation [8]. However, scATACseq subtle differences to iLSC suggest late-epi may have a more superficial/supra basal identity, with lower *KRT14* and higher *MUC16* chromatin accessibility signals. Therefore it is unclear if these cells are on track towards iLSC or already off track to a close but non-LSC fate. Further elucidation would be relevant to access the risk of this cell state for future cell therapies. To note, PAX6 was not detected in this GRN analysis due to its low expression at the mRNA level in iLSCs, although it is expressed in *in vivo* LSC and had a high influence score in GRNs derived from *in vivo* LSC scRNA-seq and scATAC-seq datasets [30]. Nevertheless, PAX6 protein expression was previously detected in iLSC [8].

A number of diverse TFs with known function in early epithelial and mesenchymal development showed high influence scores shared in several cell states. Generally, these TFs split into two groups: i) TFs mainly shared between ML2/3, such as FOXC2, NR2F1, PITX2, pointing to a neural crest identity of these states; and ii) those between ML1, EE, late-epi and iLSC, including MEIS1, GATA2/3 and TFAP2A. This TF influence pattern is consistent with WOT analysis showing that ML1/EE were on-track towards iLSC, and ML2/3 were off-track populations. Interestingly, among TFs with high influence scores in ML2, FOXC1 also showed a high influence score in iLSCs, which is consistent with previous findings that FOXC1 is required for LSC identity [35]. Although our data showed ML2/3 being likely off-track cell states, we cannot completely exclude the possibility that ML2/3 and iLSC states can influence each other in the differentiation trajectory, which warrants further investigation [36, 37].

As persistent PSC-like cells in the final iLSC culture would significantly limit the applicability of iLSC in cell therapies, it is of importance to identify the driving mechanisms of this PSC-like cell population. Previous attempts to identify TFs driving the differences did not result in significant outcome [8].Using the GRN approach, however, we found that PAX5, HOXB2 and SP8 are TFs with high influence scores and therefore are potentially key in shaping the PSC-like cell identity, distinct from the PSC state. GRN analysis further suggested that these TFs regulate each other within this state where their co-regulation may be important for a PSC-like–specific transcriptional program. The greater effectiveness of this GRN approach in predicting cell state-determining TFs may be attributed to its incorporation of genomic accessibility signals around these genes in a cell state-specific manner, even though their expression was undetectable in PSC at day 0 (undifferentiated state). Indeed, higher scATAC-seq signals near these genes were detected in PSC-like than in PSC. During differentiation, expression of *PAX5*, *HOXB2* and *SP8* were significantly higher in the hiPSC1, as compared to that in hESC, especially at day 24. This argues that hiPSC1 has the propensity to have higher expression of these genes than the ESC line, and thus inducing the PSC-like cell state. Although it would be tempting to speculate that the observed variability could reflect line-specific differentiation propensities, more differentiation rounds of these (and more) PSC lines would be required to confirm this statement. Accordingly, we have recently shown that not just line-specific but also batch to batch variability can occur within independent differentiation rounds of the same PSC line [7]. Pinpointing the source of this variability within PSC lines would be game-changing to reduce off target cell states and generate high and homogenous yields of iLSC cell populations.

Both HOXB2 [38] and PAX5 [39] have been shown to induce PI3K signaling, consistent with our earlier finding that PI3K pathway enrichment scores are higher in PSC-like cells than in PSCs [8]. SP8 can be induced by Wnt/β-catenin signaling and then activate a subset of Wnt target genes in mouse ESC [40]. Additionally, PI3K and WNT pathways are shown to enhance each other by a feed-forward loop [41], and it was also found to be elevated in PSC-like cells relative to PSCs [8]. These signaling pathways, together with the key TFs, may play a role in the line-dependent differentiation efficiency. Thus, PI3K and WNT pathways may be attractive targets to eliminate or reduce the PSC-like cell population for differentiation optimization strategies. In line with this concept, recent reports showed that differentiation-incompetent iPSC lines exhibit increased WNT pathway activation and that chemical modulation of the chromatin could decrease cell line-specific variability in neuronal differentiation [26]. It would be of great benefit in the stem cell field to further investigate the feasibility of manipulating these pathways, together with the chromatin landscape, to enhance the differentiation efficiency and consistency by reducing line-dependent variability.

In summary, we presented an integrative single-cell transcriptomic and chromatin accessibility map of iLSC differentiation, revealing distinct molecular trajectories of diverse cell states. Our findings demonstrated that enhancer-mediated regulation plays a major role in cell state specification. By integrating gene expression and chromatin accessibility within gene regulatory networks, we identified key TFs that govern these cell states. This integrative approach seems to be more sensitive to detect cell state driving TFs than gene expression data alone. Our study also provided insights into potential strategies to enhance differentiation efficiency and reduce line-dependent viability, contributing to the development of more consistent and effective corneal stem cell therapies.

## Materials and methods

### PSC differentiation into iLSC

Human ESC line Regea08/017 (“hESC”) and human iPSC line WT001.TAU.bB2 (“hiPSC1”) were previously generated and characterised in house [42].Culture conditions and differentiation into iLSC was performed as previously described by Vattulainen et al [8, 43, 44]. All cell culture experiments were performed at Tampere University, Faculty of Medicine and Health Technology. The faculty holds approval from the Finnish Medicines Agency (Fimea; Dnro FIMEA/2020/003758) to conduct research involving human embryos. In addition, the Regional Ethics Committee of the Expert Responsibility Area of Tampere University Hospital has provided supportive statements permitting the derivation, culture, and differentiation of human embryonic stem cell (hESC) lines (R05116), as well as the establishment and use of human induced pluripotent stem cell (hiPSC) lines for ophthalmic research (R16116). No new cell lines were generated for this study.

### Single-cell ATAC sequencing library preparation

At differentiation days 0, 11 and 25, cells were detached and live cells were counted following our previous study [8]. The plate-based scATACseq protocol was adapted from published protocols [45, 46]. Briefly, cells were washed twice with cold DPBS and divided into batches of 50 000 live cells for tagmentation, using 5 µL of in house produced Tn5, 45 μl tagmentation mix (20mM Tris pH 7.6, 10mM magnesium chloride, 20% dimethylformamide) and 0.25 μl Digitonin, followed by incubation at 37°C for 30min at 800rpm. Tagmentation reaction was stopped by addition of 50 μl of Tagmentation Stop Buffer (10mM Tris-HCL PH 7.8, 20mM EDTA) and kept on ice for 10 min. DAPI-positive single cells were sorted onto previously prepared 384-well plates containing lysis buffer (100mM Tris-HCL pH 8.0, 100mM NaCL, 40 μg/mL Proteinase K and 0.4% SDS) and 5 μM of both i5 and i7 index primers, using FACSAria fusion cell sorter (BD Biosciences) and stored at −80°C until further use. A minimum of two 384-well plates per differentiation round per timepoint were collected. Library amplification was performed with 2x Kappa Hifi Hot start ReadyMix (Roche) with the following cycle conditions: 72°C for 5 min, 98°C for 5 min and 20 cycles of 98°C for 10s, 63°C for 30s, 72°C for 20s. Cells from each plate were pooled and DNA purification was done with Qiagen minElute PCR Purification Kit (Qiagen) with final elution volume of 20 μL. Size selection was done with 1.5x Agencourt AMPure XP beads (Beckman Coulter) and samples ran in BioAnalyzer High Sensitivity DNA Kit (Agilent Technologies) for quality control. Samples were sequenced on a NextSeq 500 (Illumina) for 30 million reads/plate.

Data in libraries were processed with seq2science to generate trimmed fastq files per single well. Data from single wells were joined with a custom python script and a snakemake pipeline was used to remove duplicate reads with Picard[47] and generate bam files for each 384 well plate.

### Data availability

Previously published scRNA-seq was downloaded from Zenodo [48]. scATACseq data was deposited on the Gene Expression Omnibus with the identifier GSE313398, The full documentation of computational workflows, including scripts and snakemake pipelines, is available at the GitHub repository: https://github.com/Arts-of-coding/Characterizing-iPSC-derived-LSC-differentiation-through-integrative-single-cell-omic-approaches.

### scRNA-seq optimal transport analysis

Waddington-OT (WOT) [11] (version 1.0.8) was used according to provided documentation on scRNA-seq data in a Python software environment (version 3.7.12) with Scanpy (version 1.9.3) [49]. In short, transport maps were generated for each cell line across the three timepoints, by linking timepoints 1 (day 0) and 2 (day10/11) as well as 2 and 3 (day 24). From these maps, comprehensive networks were generated. Similarly, WOT cell fate scores were calculated by linking timepoints in a similar fashion. Additionally, networkx (version 2.6.3) [50] was used to generate overview network graphs of these transport maps.

### scATAC-seq analysis

SnapATAC2 (version 2.8.0) [16] was used to process scATAC-seq data in a Python software environment (version 3.10.12). First, fragment files were generated through importing processed bam files. Next, cells were filtered with a minimum fragment number of 5000, transcription start site enrichment (tsse) scores higher than 2 and a maximum number of counts smaller than 100.000. Following, the top 250.000 features were selected and scrublet [51] was used to remove doublets. Dimensionality reduction was performed with spectral (Laplacian Eigenmaps), according to provided documentation. Data was integrated with Harmony [17] on the two experimental rounds selected as batch and was subsequently visualized with Uniform Manifold Approximation and Projection (UMAP). For visualization purposes, the y-axis of the generated UMAP was inverted, pseudo expression was calculated by generating a gene matrix and magic [52] imputation was used to impute counts for cells in close proximity.

### Genome tracks

Bam files were merged into a single bam file with samtools [53]. This merged bam file was split on cluster and cell state specific barcode csv files with cellranger-dna’s subset-bam [54]. Peak calling was performed with macs3 [55]. Bigwig files were generated with UCSC’s bedGraphToBigWig as previously described [30] and were subsequently loaded into the UCSC genome browser.

### scATAC-seq peak analysis

Peaks were defined on the cluster level with Snapatac2’s merge peaks. Unique peaks were defined as being present only in one cluster or cell state and not in any of the others. A promoter region bed file was generated with bedtools flank [56] using the annotated genome together with a 2kb size before the gene start. Next, this file was used to allocate promoter and non-promoter (enhancer) peaks for all identified peaks and for unique peaks. The enrichment (odds-ratio) for enhancer peaks over promoter peaks, for comparing unique peaks against all peaks, was calculated with Fisher’s exact test using scipy (version 1.13.1) [57].

### Enhancer to gene signal calculation

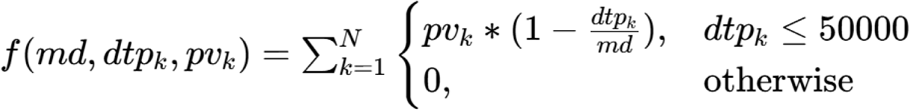

For each gene, signals of enhancers were weighed and summed based on their genomic distance [1]. Where *dtp_k_*is the distance of a counted enhancer to a gene of interest (GOI), *pv_k_*is the raw peak count value for each enhancer and *md* is the maximum genomic distance to a gene (50 kilobases). Enhancer to gene signals were scaled to the respective cell state by dividing the total number of cells for that cell state with the lowest number of cells from all cell states.

### scRNA-seq and scATAC-seq integration and prediction

scRNA-seq data was integrated with scATAC-seq data by scANVI [20] as described by Snapatac2 documentation. In short, a scVI model was constructed with the top 5000 highly variable genes with 2 layers and 30 latent layers as model parameters. The model was trained for 500 epochs with early stopping enabled. Subsequently, a scANVI model was constructed from the pre-trained scVI model and scRNA-seq predictions were made on scATAC-seq data. Lastly, the fraction of predicted cell state identity in scATAC-seq based on scRNA-seq was calculated for each identified cluster (Supplementary Table S3).

### Gene regulatory network analysis

scANANSE was performed on multiple single-cell comparisons with AnanseScanpy [15] as described in Arts et al,. 2024 [30]. First, GRNs were generated by using all identified peaks and all unique peaks from Snapatac2 for each cell state. Underlying scATAC-seq data was used in multiple networks if the scATAC-seq cell state was annotated with multiple scRNA-seq cell state names, for instance ML2/ML3 was both used for generating the meso-like-2 and the meso-like-3 networks. The first comparison was done with specific cell state networks against a network from the average of all cell states, described in the scANANSE workflow. The second comparison directly compared PSC and PSC-like networks to each other.

### Quantitative RT-qPCR analysis of PSC and iLSC

The undifferentiated PSCs as well as differentiated cells at d11 and d24 timepoints of the differentiation were analyzed for their mRNA expression with RT-qPCR. TaqMan Gene Expression Assays (Applied Biosystems, MA, USA) for *PAX5* (Hs05040337), *SP8* (Hs01941366) and *HOXB2* (Hs01911167) were used to analyze gene expression at all time points GAPDH was used as a housekeeping gene. RNeasy Mini Kit (Qiagen, Netherlands) were used for RNA isolation and cDNA was synthesized by using reverse transcription with a High-capacity cDNA kit (Applied Biosystems, United States) from the cell pellet samples collected in indicated time points. Biological replicates (n = 3) and controls were run as triplicate reactions with the ABI QuantStudio 12K Flex Real-Time PCR system (Applied Biosystems). The results were analyzed using the −□2ΔΔCt method and are presented as the fold change in gene expression normalized to GAPDH and relative to the undifferentiated controls.

All data are presented as individual values. The Mann-Whitney *U* test was performed to analyze the differences between the groups using the GraphPad Prism 10 software (GraphPad Software Inc.). Differences were considered statistically significant when *P*□≤ 0.05. Number of samples (*n*) is constituted of iLSC differentiation batches, each of individual cell batches being considered as biological replicates.

## Acknowledgements

The authors thank Outi Melin and Hanna Pekkanen (Tampere University) for cell production and technical assistance; Marc Eleveld (Radboud University Medical Centre) and Laura Wingens (Radboud University) for assistance in scATAC plate preparation; and Marijke Baltissen and Sybren Rinzema (Radboud University) for sequencing technical assistance and data management. We also acknowledge the Biocenter Finland (BF), Tampere Imaging Facility (TIF) and Tampere Flow Cytometry Facility for their service.

This project was funded by the Netherlands Organisation for Health Research and Development (ZonMW) projects ZonMw Open (09120012010039) to H.Z., J.A.A., D.L.C., Off Road (04510012210043) to D.L.C and PSIDER TOP (10250052410019) to D.L.C., H.Z; COST Action CA18116 Aniridia-Net for Short-Term Scientific Mission (STSM) grants to D.L.C., I.B; and Research Council of Finland (338988; 370564) and the Sigrid Jusélius Foundation to H.S.

## Author contributions

Conceptualization: J.A.A, D.L.C, M.V., F.B., H.S., H.Z.; methodology: J.A.A., D.L.C., S.H., M.V., J.G.A.S., I.B., D.F.B.; software: J.A.A., J.G.A.S., D.F.B., F.B.; formal analysis: J.A.A., D.L.C., S.H.; data curation: J.A.A., H.Z.; investigation: J.A.A., D.L.C., S.H., M.V.; resources: H.S., H.Z.; writing – original draft: J.A.A., D.L.C. and H.Z.; writing – review and editing: all authors

