## Supplementary Data for "Mapping heterogeneity drivers in human PSC-derived limbal stem cell differentiation through single-cell multi-modal analysis"

**SUPPLEMENTARY FIGURES**

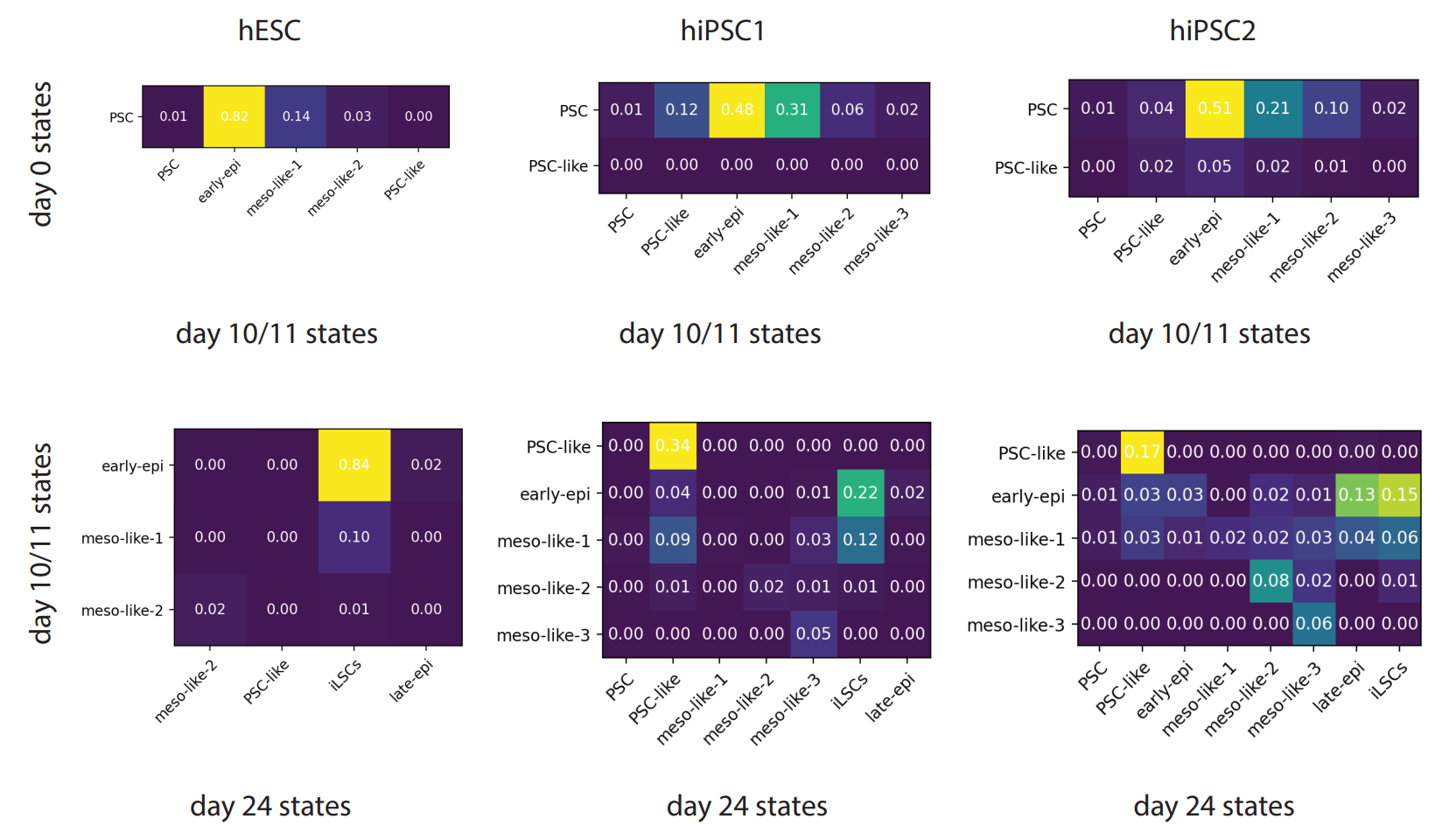

***Supplementary Figure 1: Waddington-OT transition tables from time points of iLSC differentiation.*** *Transition tables depict predicted cell fractions transitioning into each other from day 0 to day 10/11 as well as day 10/11 to day 24 for the three cell lines hESC, hiPSC1 and hiPSC2. Higher percentages were highlighted with brighter colours.*

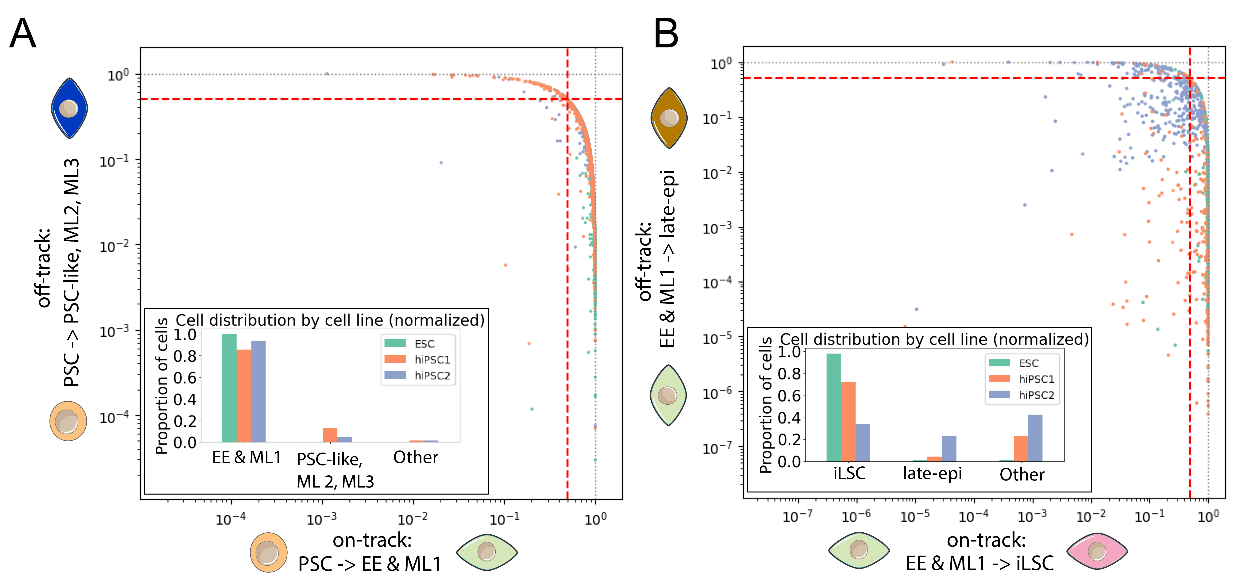

***Supplementary Figure 2: Waddington-OT cell fate scores of differentiating cell states.*** *(scale 0-1) for PSC (day 0) that differentiate into EE and ML1 (day10/11) against PSC-like, ML2, ML3 (A), and EE and ML1 into iLSC against late-epi (B). Bar plot quantifications of WOT cell fate scores of PSC (E) and EE/ML1 (F) differentiating into cell states at subsequent timepoints. Cells towards either cell state(s) indicated on the X and Y axis are divided by the cell fate score 0.5 into three categories (the dotted orange line): two categories with scores >0.5 and the category other that refers to cells with cell fate scores <0.5.*

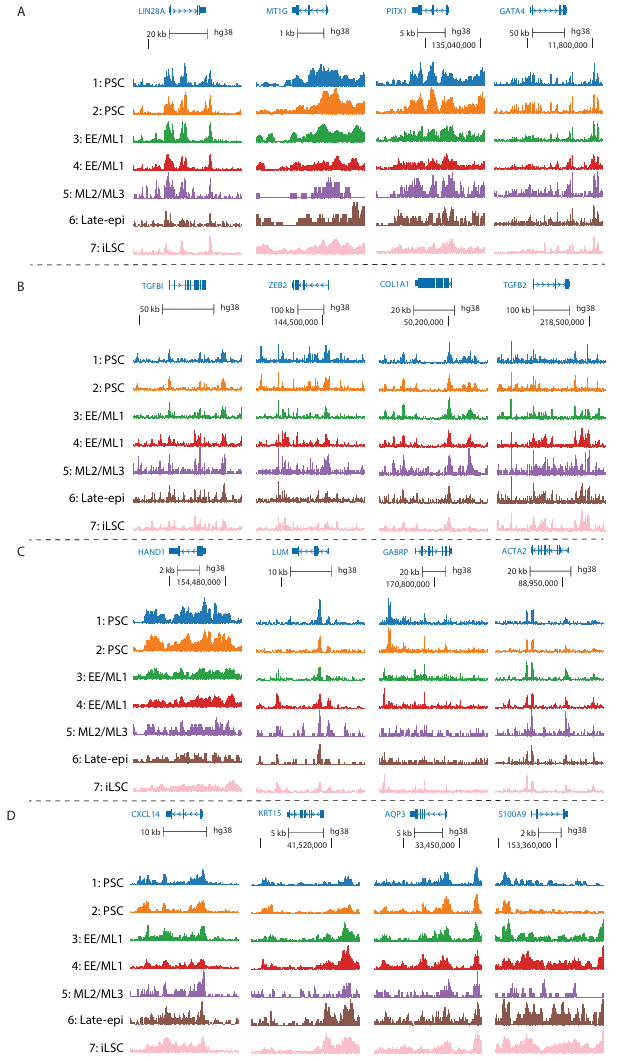

***Supplementary Figure 3:*** *UCSC genome browser tracks (A-D) of LIN28A, MT1G, PITX1 and GATA4 (A), TGFBI, ZEB2, COL1A1 and TFGB2 (B), HAND1, LUM, GARBP and ACTA2 (C) and CXCL14, KRT15, AQP3 and S100A9 (D).*

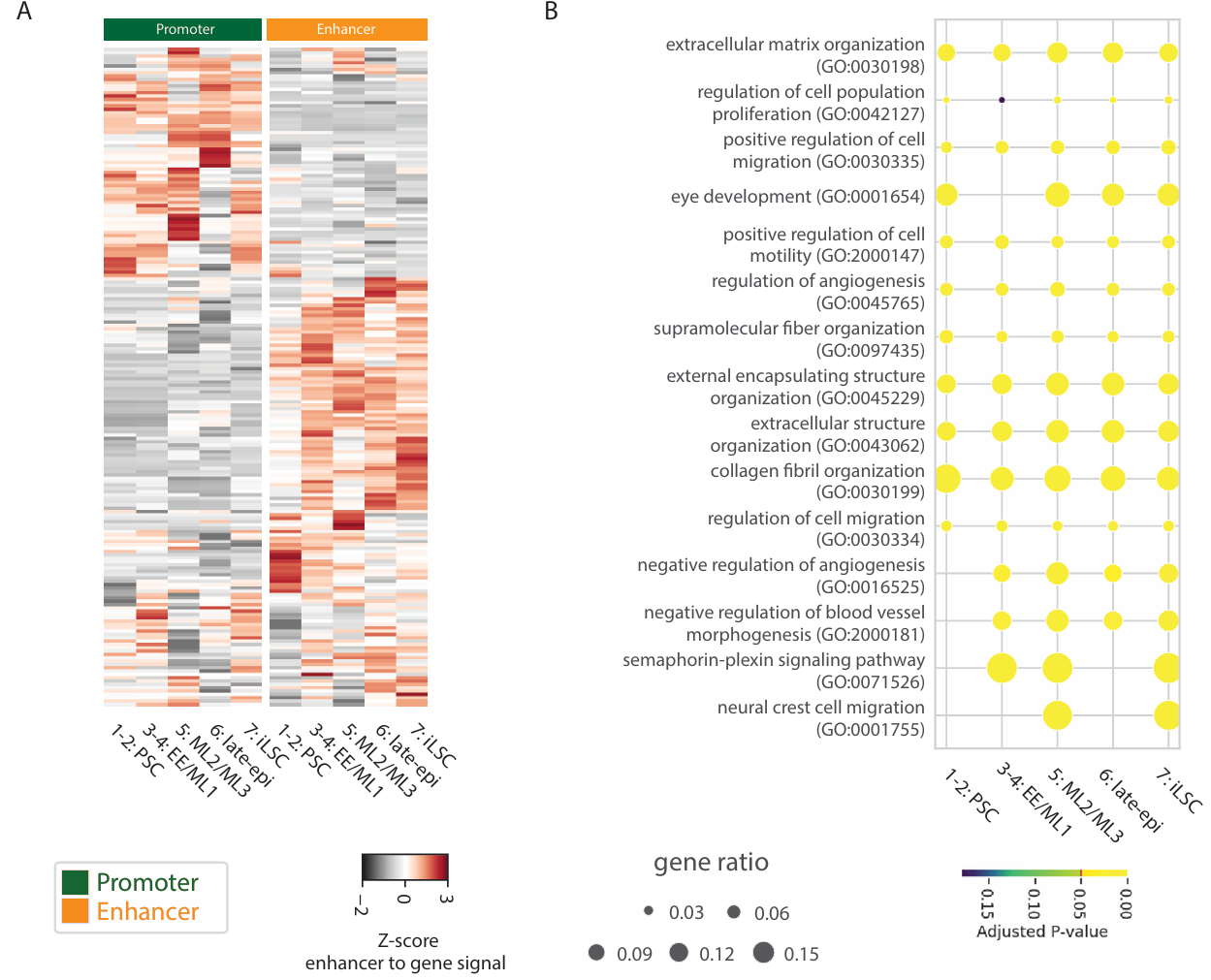

***Supplementary Figure 4: Biological processes identified for all identified peaks linked to highly variable genes.*** *A) Heatmaps of scATAC-seq depicting Z-scores from peaks linked to previously defined highly variable genes in scRNA-seq (log2FC>1.5 and p-adjusted value <0.05). The top 5 HVGs with highest peak signal at enhancers are located on the right of the plot. B) Top 10 merged GO-terms on enhancer-to-genes signal of previously identified HVGs (log2fc>1.5, padj < 0.05) for unique peaks. Dot size indicates the ratio of genes found for a specific GO-term. The color of the dot indicates the p-adjusted value.*

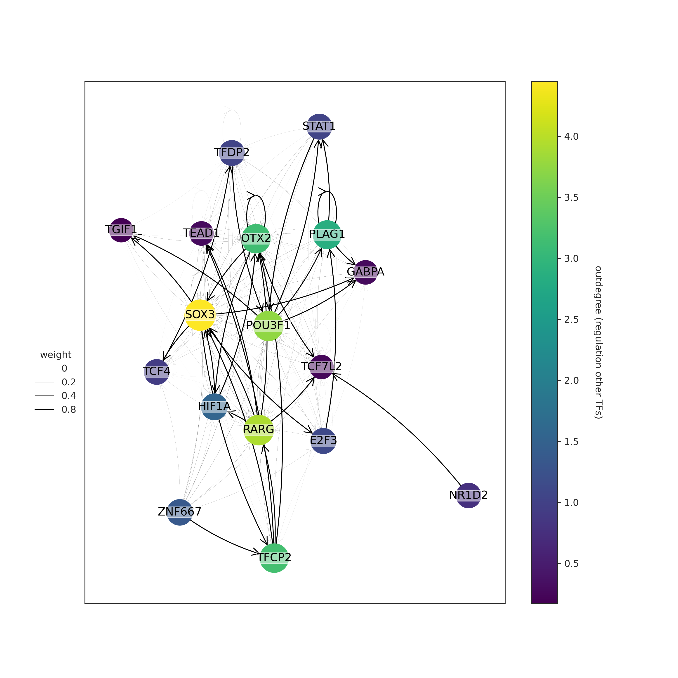

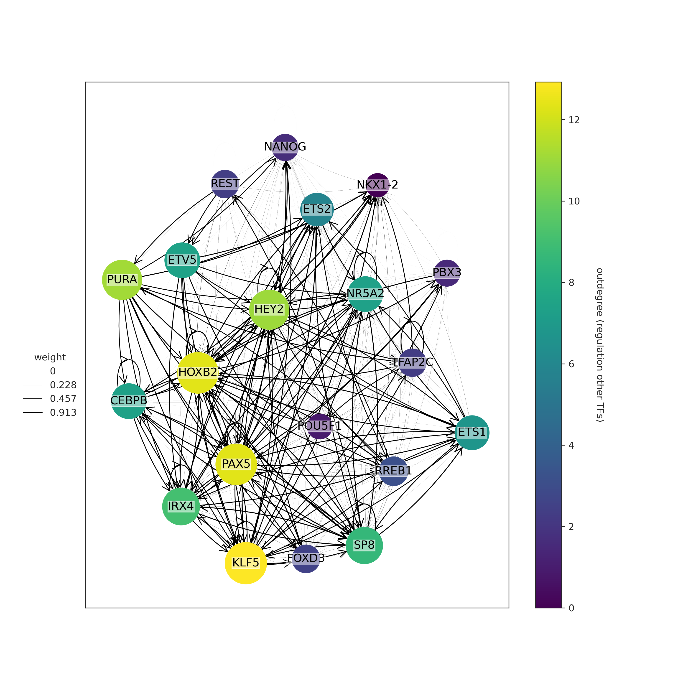

***Supplementary Figure 5:*** *scANANSE’s interacting TF networks using differential TFs in the PSC/PSC-like comparison. (A) PSC over PSC-like; (B) PSC-like over PSC*

**SUPPLEMENTARY TABLES**

***Supplementary Table 1****: Distribution of the absolute number of cells across cell lines in scRNA-seq across time points.*

| **scRNA-seq cell state** | **Day0 ESC** | **Day 0 hiPSC1** | **Day 0 hiPSC2** | **Day 10/11 ESC** | **Day 10/11 hiPSC1** | **Day 10/11 hiPSC2** | **Day 24 ESC** | **Day 24 hiPSC1** | **Day 24 hiPSC2** |
| --- | --- | --- | --- | --- | --- | --- | --- | --- | --- |
| PSC | 735 | 741 | 637 | 3 | 5 | 5 | 0 | 2 | 7 |
| PSC-like | 0 | 2 | 87 | 1 | 39 | 34 | 1 | 151 | 95 |
| early-epi | 0 | 0 | 1 | 429 | 164 | 283 | 0 | 0 | 18 |
| meso-like-1 | 0 | 0 | 0 | 74 | 104 | 118 | 0 | 1 | 8 |
| meso-like-2 | 0 | 0 | 0 | 17 | 19 | 54 | 16 | 7 | 51 |
| meso-like-3 | 0 | 0 | 0 | 0 | 7 | 12 | 0 | 33 | 53 |
| late-epi | 0 | 0 | 0 | 0 | 0 | 0 | 10 | 6 | 75 |
| iLSC | 0 | 0 | 0 | 0 | 0 | 0 | 539 | 109 | 87 |
| Total cells | 735 | 743 | 725 | 524 | 338 | 506 | 566 | 309 | 394 |

***Supplementary Table 2.*** *Fraction tables (range 0-1) of WOT transitions depicting the fraction of the total of cells of a cell line transitioning into a certain category (above or below cell fate score 0.5) used in Supplementary Figure 2. WOT cell fate scores (scale 0-1) of cells transitioning from PSC to day 11 cell states (top) and from EE/ML1 (day 11) to day 24 cell states (bottom). “Other” refers to cells with cell fate score lower than 0.5.*

| **PSC to:** | **ESC** | **hiPSC1** | **hiPSC2** |
| --- | --- | --- | --- |
| EE, ML1 | 0.99 | 0.85 | 0.93 |
| ML2, ML3, PSC-like | 0.001 | 0.13 | 0.05 |
| Other | 0.001 | 0.02 | 0.05 |
| **EE/ML1 to:** |  |  |  |
| iLSC | 0.98 | 0.72 | 0.34 |
| Late-epi | 0.01 | 0.04 | 0.23 |
| Other | 0.01 | 0.24 | 0.43 |

***Supplementary Table 3****: Total cell counts across scATAC-seq clusters and percentages from figure 3 B-D.*

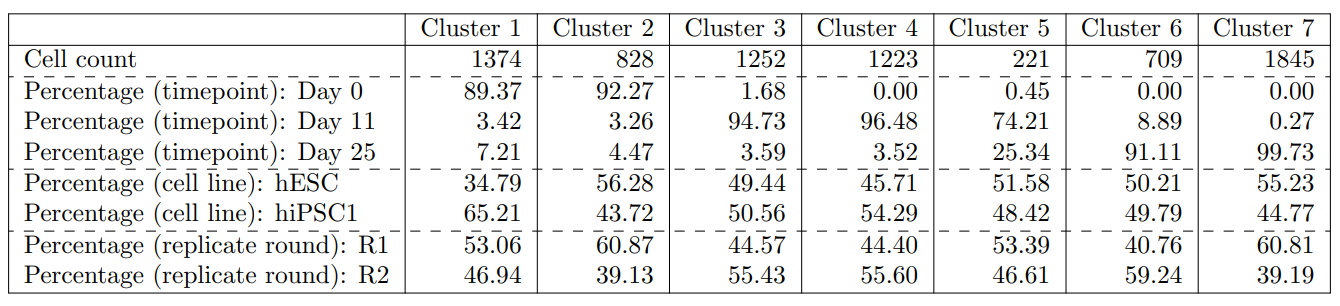

***Supplementary Table 4****: Cluster proportions and permutation analysis from scATAC-seq data, separated by PSC line.*

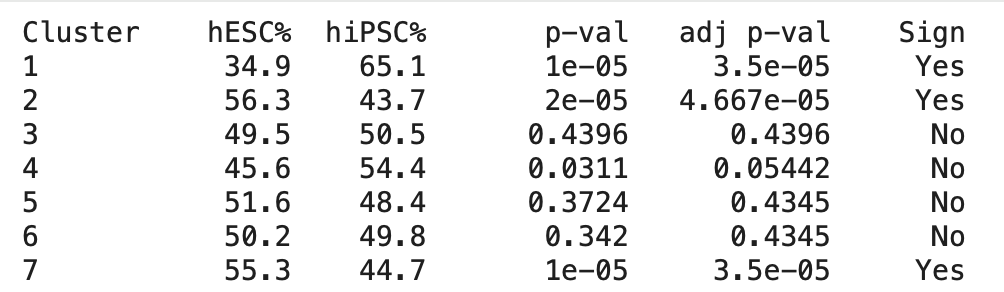

***Supplementary Table 5****: Cell fractions (range 0-1) of scRNA-seq predicted labels of cells in scATAC-seq clusters, calculated with scANVI.*

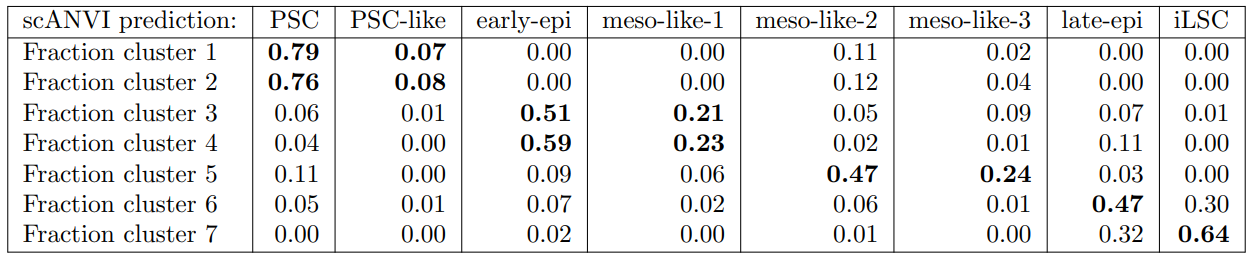

***Supplementary Table 6****: Percentages of promoter and enhancer regions from all identified peaks in each cell state.*

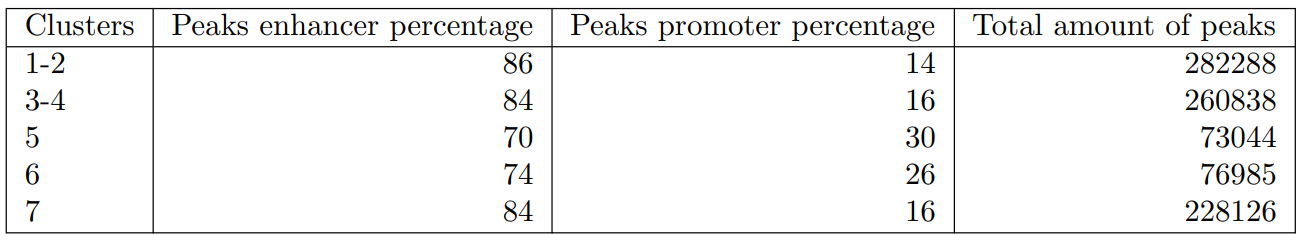

***Supplementary Table 7****: Percentages of promoter and enhancer regions from unique identified peaks in each cell state.*
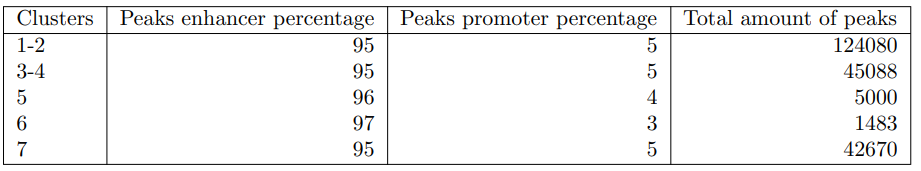

***Supplementary Table 8****: TFs with scANANSE’s influence scores higher than 0.6 (scale 0-1) for the iLSC network constructed from unique peaks compared to the average network (Figure 4B).*

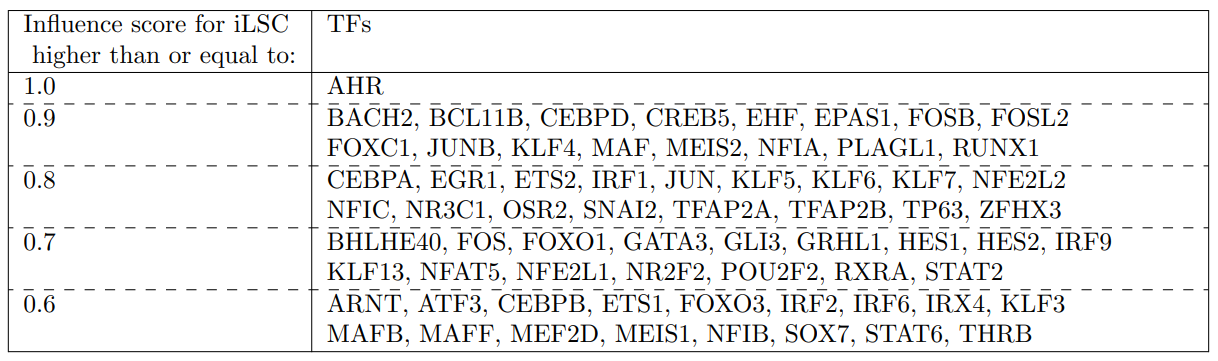

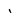
